# Genome-wide analysis resolves the radiation of New Zealand’s freshwater *Galaxias vulgaris* complex and reveals a candidate species obscured by mitochondrial capture

**DOI:** 10.1101/2022.07.04.498769

**Authors:** Ciaran S.M. Campbell, Ludovic Dutoit, Tania M. King, Dave Craw, Christopher P. Burridge, Graham P. Wallis, Jonathan M. Waters

## Abstract

Freshwater fish radiations are often characterized by multiple closely-related species in close proximity, which can lead to introgression and associated discordance of mitochondrial and nuclear characterizations of species diversity. As a case in point, single locus nuclear versus mitochondrial analyses of New Zealand’s stream-resident *Galaxias vulgaris* complex have yielded conflicting phylogenies. We generate and analyze a genome-wide data set comprising 52,352 SNPs across 187 *Galaxias* specimens to resolve the phylogeny of this recent fish radiation. We conduct phylogenetic, PCA, STRUCTURE, and ABBA-BABA analyses to evaluate the evolutionary relationships of lineages in the context of natural and anthropogenic river drainage alterations. In addition to the 11 previously recognized stream-resident lineages, genome-wide data reveal a twelfth candidate species (*G*. ‘Pomahaka’), apparently obscured by introgressive mitochondrial capture. We identify additional examples of mito-nuclear discordance and putative mitochondrial capture, likely mediated by geological and anthropogenic modification of drainage boundaries. Our study highlights the need for genome-wide approaches for delimiting freshwater biodiversity. Genetic data also reveal the influence of drainage history on freshwater biodiversity, including the rapid divergence of recently fragmented fish populations, and the conservation genetic risks of anthropogenic translocations events.

## 1. INTRODUCTION

As anthropogenic pressures on freshwater ecosystems increase, reliable delimitation of species is increasingly important for the conservation of freshwater biological diversity (Allendorf et al., 2022; Closs et al., 2016; Olden et al 2010). To this end, the analysis of genome-wide data is transforming our understanding of freshwater biodiversity across a range of spatial and evolutionary scales (Melo et al., 2021; Ronco et al., 2021). Indeed, freshwater fish assemblages often exhibit astonishing levels of diversity considering the restricted nature of habitats they occupy (Adams et al., 2014; Raadik et al., 2014; Ronco et al., 2021; Shelley et al., 2018). This high diversity may be partly explained by divergence of isolated freshwater populations in the absence of marine links through salinity tolerance or migratory (diadromous) life history (Burridge & Waters, 2020; Delgado et al., 2020; Ward et al., 1994; Waters et al., 2020a). In addition to these effects of geographic isolation *per se* (Waters et al., 2020a), dynamic geological processes (Craw et al., 2015; Waters et al., 2001a, 2020b) and rapid adaptive shifts can also contribute substantially to freshwater biological diversification (Barluenga et al., 2006; Melo et al., 2021; Ronco et al., 2021).

Freshwater-limited fish radiations are often characterized by multiple closely-related species in close proximity, and hence are particularly prone to hybridization and introgression (Irisarri et al., 2018; MacGuigan & Near, 2019; Wallis et al., 2017; Waters et al., 2010). This mixing may be exacerbated via human-driven range shifts (Blackwell et al., 2021; Esa et al., 2000), further complicating the estimation of phylogenetic relationships and systematics. Given that distinct genomic regions can vary in their propensities for introgression, freshwater lineages are particularly prone to mitonuclear discordance (Ford et al., 2019; Wallis et al., 2017). As a case in point, introgressive *mitochondrial capture* (Perea et al., 2016; Unmack et al., 2011; Willis et al., 2014) – whereby the mitochondrial DNA (mtDNA) of one lineage is completely replaced by that of another – represents the most extreme form of such discordance and has potential to severely compromise the detection and conservation of biodiversity. However, the recent application of genome-wide data (e.g. Elshire et al., 2011) to freshwater systematics has begun to resolve such discordance (Buckley et al., 2018; Perea et al., 2016; Unmack et al., 2017), enhancing the recognition and conservation of freshwater biodiversity.

The widespread ‘Gondwanan’ galaxiid fishes represent a key component of the Southern Hemisphere’s freshwater fish fauna, comprising a combination of diadromous and freshwater-limited taxa (Burridge et al., 2012; McDowall, 1970, 1990). In recent decades, genetic analyses have detected substantial cryptic species diversity, particularly within various freshwater-limited galaxiid lineages, leading to the recognition of several distinct species-rich complexes (Adams et al., 2014; Allibone et al., 1996; Chakona et al., 2013; Waters & Wallis, 2001a, b). These findings have led to the description of numerous new taxa (e.g. McDowall, 1997; McDowall & Wallis, 1996; Raadik, 2014). New Zealand’s (NZ’s) *Galaxias vulgaris* complex, for example, was once thought to comprise just a single stream-resident taxon (McDowall, 1970, 1990), but is now recognized as comprising 11 lineages (including six formally described species along with five additional undescribed lineages; Figure 1; Burridge et al., 2012; Waters et al., 2010). Many of these distinctive freshwater lineages differ genetically, morphologically and ecologically (e.g. Allibone & Townsend, 1997; Crow et al., 2009), with distinctive ‘flathead’ versus ‘roundhead’ lineages (Figure 1) often co-occurring, facilitating tests for reproductive isolation (e.g. Crow et al., 2009; McDowall & Wallis, 1996; Waters et al., 2001b). In the context of widespread habitat alteration and the introduction of salmonid predators, conserving this endemic freshwater biodiversity presents a major conservation challenge (Dunn et al., 2018; McDowall, 2006).

**FIGURE 1.**
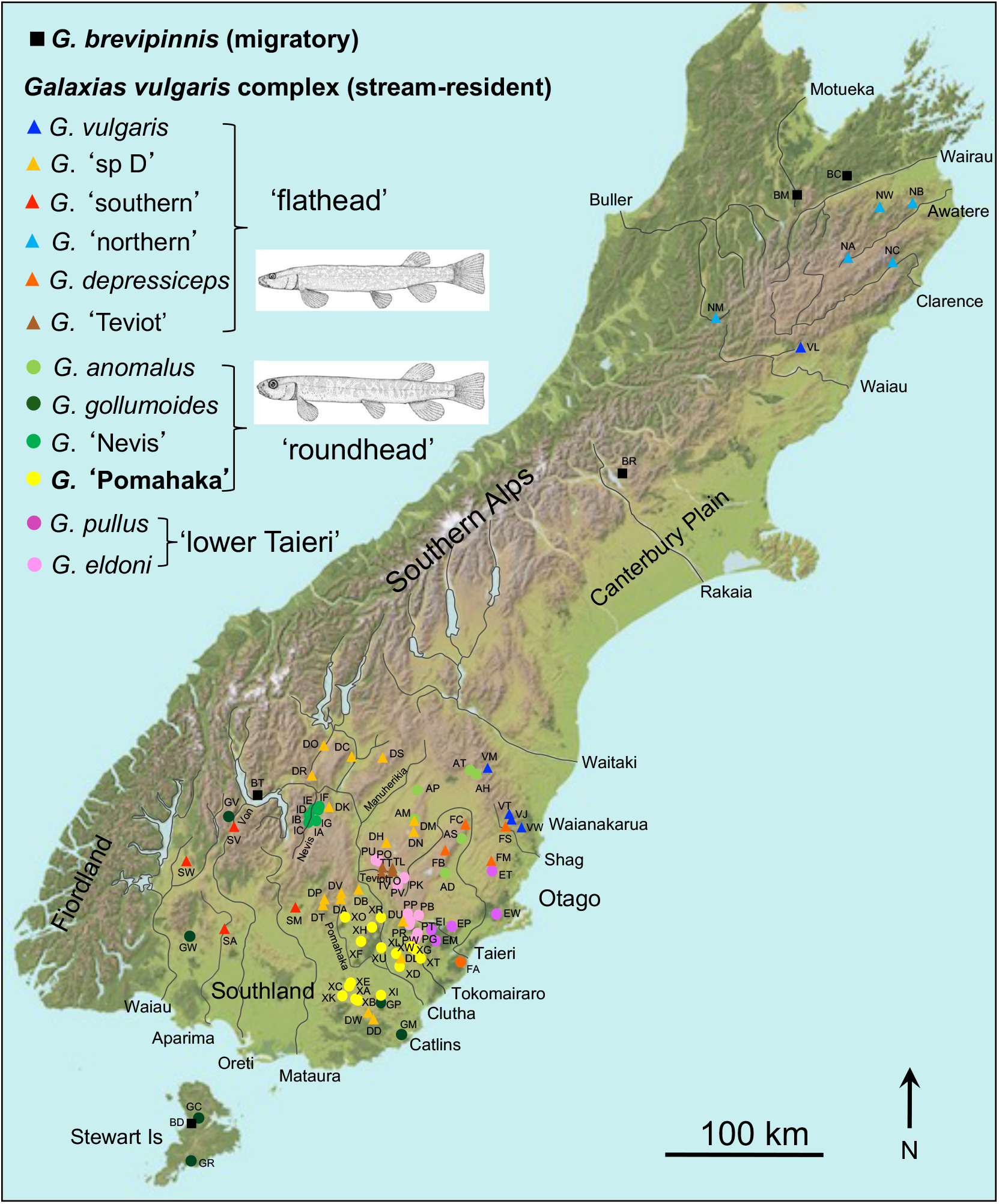
New Zealand sampling localities for *Galaxias* specimens included in genome-wide analyses. Sampling locality/lineage codes are included in Table S1. The newly-recognised lower Clutha ‘roundhead’ lineage *Galaxias* ‘Pomahaka’ (bold; yellow) detected using genome-wide analysis carries mtDNA identical to the unrelated but co-distributed Clutha ‘flathead’ *Galaxias* ‘sp D’ (orange), suggesting introgressive mitochondrial capture. Representative ‘flathead’ (triangles) and ‘roundhead’ (circles) morphotypes are indicated (from McDowall & Wallis, 1996).

New Zealand’s *G. vulgaris* complex shares a common ancestry with the widespread diadromous taxon *G. brevipinnis* (shared by Australia and NZ; McDowall, 1990). This distinctive freshwater-limited assemblage was initially suggested to have evolved via numerous independent transitions from diadromous to stream-resident life-history (Allibone & Wallis, 1993; Waters & Wallis, 2001b; Figure 2), as is the case with several freshwater-limited fish complexes elsewhere (Colosimo et al., 2005; Delgado et al., 2019; 2020; Fang et al., 2020; Veale et al., 2017). However, a subsequent multilocus phylogeny found the freshwater-limited *G. vulgaris* complex to be monophyletic (Figure 2), and thus concluded that the radiation can be explained by a single loss of diadromy (Waters et al., 2010). Multidisciplinary studies combining geological and biological data (predominantly mtDNA) suggest that geological processes such as mountain building and river capture (Burridge et al., 2008a; Craw et al., 2015) have been highly influential in the diversification and spread of these stream-resident lineages (e.g. Waters et al., 2001a, 2020b).

**FIGURE 2.**
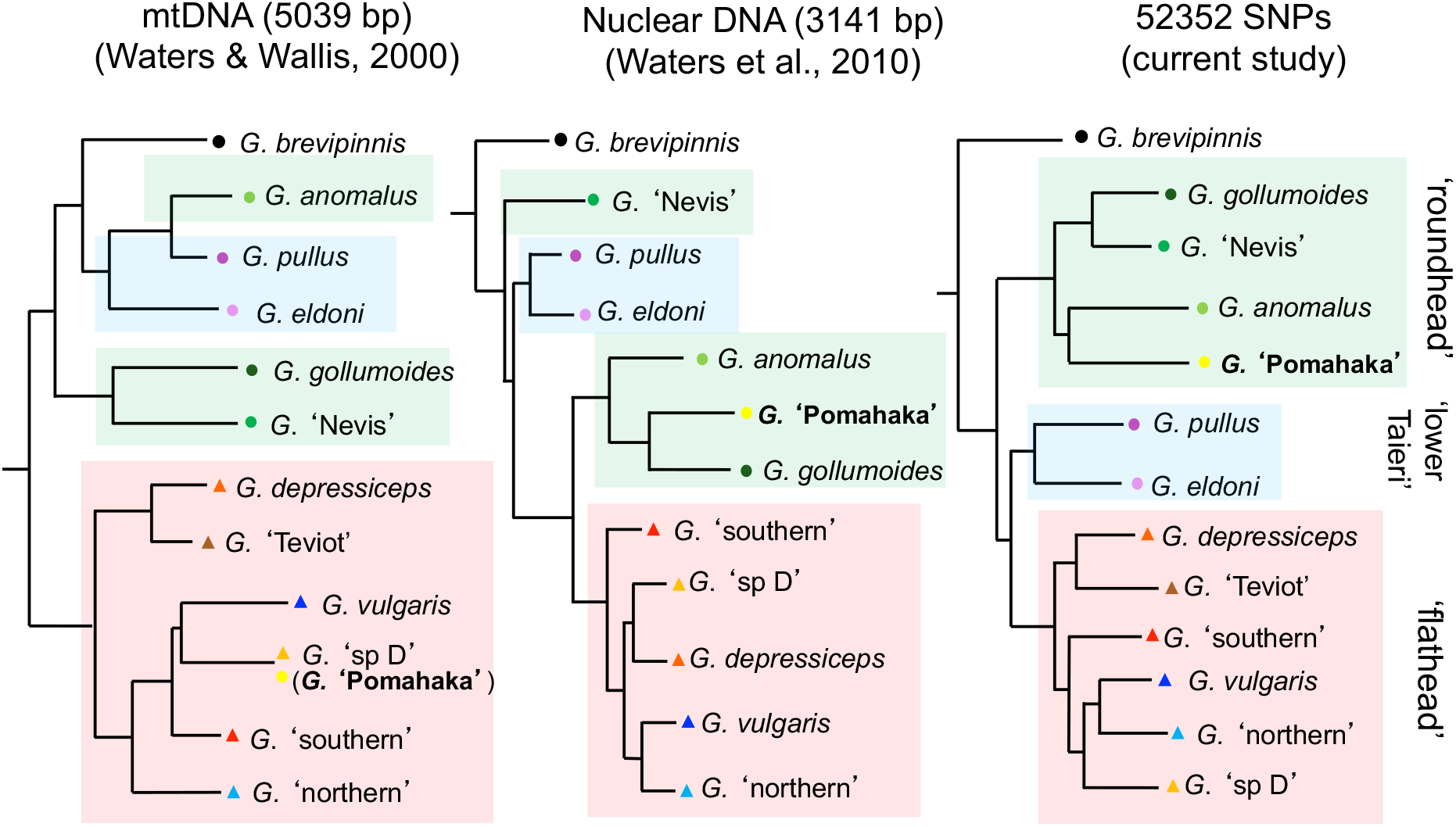
Discordant mito-nuclear phylogenetic relationships of New Zealand’s *Galaxias vulgaris* complex based on mtDNA (left; redrawn from Waters & Wallis, 2001a) versus concatenated nuclear sequence data (centre; redrawn from Waters et al., 2010) and genome-wide SNPs (right; full details in Figure 3). Distinctive ‘flathead’ (triangles) versus ‘roundhead’ (circles) assemblages broadly characterized by morphological, life-history and ecological variation (e.g. Crow et al., 2009; Jones & Closs, 2016; Jones et al., 2016; McDowall & Wallis, 1996; McDowall, 1997) are highlighted by coloured boxes. The newly recognized ‘roundhead’ *Galaxias* ‘Pomahaka’ (bold) carries mtDNA identical to that of the sympatric ‘flathead’ *Galaxias* ‘sp D’ (left), but these two lineages are otherwise phylogenetically unrelated (centre; right). Similarly, *Galaxias anomalus* has strong mtDNA similarity with *Galaxias pullus* (left), but these two taxa are otherwise phylogenomically unrelated to each other (centre; right).

In biogeographic terms, the diversity of the *G. vulgaris* complex is centred on the Otago region of southern South Island (Figure 1), with the Taieri and Clutha river systems each housing numerous taxa (Waters et al., 2001a; Waters et al., 2010; Waters et al., 2015a). In addition to the 11 currently recognized lineages (Dunn et al., 2018), a preliminary multi-locus sequence analysis (Waters et al., 2010) detected an additional nuclear DNA lineage that remained cryptic at the mtDNA level (Burridge et al., 2007; Waters et al., 2001a). Specifically, a few individuals sampled from the Pomahaka R (a major tributary of Otago’s Clutha system; Figure 1) have yielded a distinctive nuclear DNA clade (which we here provisionally name *G*. ‘Pomahaka’), apparently phylogenetically affiliated to ‘roundhead’ taxa *G. anomalus* and *G. gollumoides* (Waters et al., 2010; Figure 2). By contrast, extensive mtDNA analyses from this region have consistently allocated all Pomahaka *Galaxias* samples (and most other lower Clutha samples) to the widespread Clutha ‘flathead’ lineage *G*. ‘sp D’ (Burridge et al., 2007; Waters et al., 2001a). Interestingly, subsequent morphological assessment has suggested the presence of both ‘roundhead’ and ‘flathead’ morphotypes within the Pomahaka system (D. Jack, pers. comm.). However, extensive mtDNA sequencing throughout the Pomahaka system has continued to yield just a single mitochondrial clade, attributable to the widespread *G*. ‘sp D’ (Waters et al., 2015b).

Here we use Genotyping-by-Sequencing (GBS; Baird et al., 2008; Elshire et al., 2008) to undertake genome-wide analyses of NZ’s species-rich *G. vulgaris* complex to fully characterize its diversity and infer the role of river-drainage alteration in its formation. In particular, we test the hypothesis that the lower Clutha system contains unrecognized species diversity. We use this system as a model for assessing the effects of introgressive mitochondrial capture on freshwater fish systematics and conservation. Our study also tests for phylogeographic divergence among recently fragmented fish populations, and assesses the genetic implications of anthropogenic drainage modification.

## 2. METHODS

### 2.1 Sampling and library preparation

We retrieved 227 ethanol-preserved *Galaxias* specimens from the University of Otago galaxiid fish collection where they are stored at -20°C. The vast majority of these samples had been collected by NZ’s Department of Conservation over the last two decades. We typically selected two specimens per locality, incorporating multiple locations for each of 11 recognized *G. vulgaris* complex lineages (Dunn et al., 2018; Waters et al., 2010), along with NZ and Australian representatives of the diadromous sister taxon *G. brevipinnis*. In addition to representing recognized stream-resident diversity, our sampling effort focused particularly on the Pomahaka R and surrounding catchments of the lower Clutha system (southern South Island), where previous studies have detected nuclear genetic diversity not evident using mitochondrial markers (Waters et al., 2010).

High-molecular weight DNA was extracted from tissue samples using a Qiagen DNeasy Blood and Tissue Kit, following the manufacturer’s protocols. The concentration and quality of eluted DNA was assessed using a DeNovix DS-11 spectrophotometer and an Invitrogen Qubit 3.0 fluorometer. DNA degradation was assessed visually using agarose gel electrophoresis. Based on concentrations, purities and visual degradation assessments, 192 samples were selected (from the original 227 samples of extracted DNA) for library preparation. These samples had DNA concentrations greater than 10ng/µL, received no purity alerts, and had suffered minimal visual degradation within the 300–600 bp range, crucial for GBS (Graham et al., 2015).

Library preparation for GBS followed methods described by Elshire et al. (2011) with modifications described in Dussex et al. (2015). The pooled library was size-selected (300– 600 bp) and sequenced at the Biomolecular Resource Facility, Australian National University using an Illumina NextSeq 500 (75 bp paired-end).

### 2.2 Bioinformatics

Quality scores and adapter content were investigated using FastQC v0.11.9 (Andrews, 2017). SNPs were called from sequence data using the *Stacks* pipeline v2.55 (Catchen et al., 2013). Briefly, the *process_radtags* module in *Stacks* was used to demultiplex sequence data, remove sequences with uncalled bases (-c) and discard low quality reads (-q). *process_radtags* removed sequences with no barcode, no cut-site, and low quality sequences.

We assembled loci without a reference genome (i.e. *de novo*) by executing the *denovo_map*.*pl* script in *Stacks*. The minimum number of reads to create a locus was set at 3 (-m), and the maximum number of pairwise differences between loci was 2 (-M). To limit numbers of SNPs in linkage disequilibrium, only the first SNP in each locus was kept for subsequent analysis (--write-single-snp). We used VCFtools v0.1.15 (Danecek et al., 2011) to exclude SNP sites with greater than 10% missing data across individuals.

### 2.3 Phylogenomic relationships and introgression

Principal Component Analysis (PCA) was undertaken in R v4.0.3 (R Core Team, 2020). The vcfR v1.10.0 package (Knaus & Grünwald, 2017) enabled R to read *Stacks* output files, and the PCA analysis was performed and visualised using pcadapt v4.3.1, poppr v2.8.5, adegenet v2.1.2 and ade4 v1.7 packages (Dray & Dufour, 2007; Jombart, 2008; Kamvar et al., 2014; Luu et al., 2017). Phylogenomic analyses were undertaken utilising RAxML (Stamatakis, 2014) under a maximum likelihood framework using a GTR-GAMMA model with 100 rapid bootstraps. FigTree v1.4.4 (Rambaut, 2009) was used to visualise and root the tree with Australian *G. brevipinnis* specified as an outgroup (see Waters et al., 2010).

Phylogenomic relationships among lineages were also assessed under a coalescent framework using SNAPP (Bryant et al., 2012), implemented in BEAST v2.6.6 (Bouckaert et al., 2019). Specifically, we reduced the dataset to four haploid sequences of 5,000 random SNPs for each candidate species as inferred from ML analyses (Table S1), and rooted the tree with Australian *G. brevipinnis*. The MCMC chain convergence was assessed using Tracer v1.7.1 and the output was visualised using DensiTree (Bouckaert, 2010). We tested for nuclear introgression among specific lineages using the ABBA-BABA approach (Green et al., 2010; Durand et al., 2011) as implemented in Dsuite v0.5. (Malinsky et al., 2021).

We used fastSTRUCTURE v1.0 (Raj et al., 2014) to assess genotypic subgroupings under a Bayesian framework. SNP data were exported and converted for fastSTRUCTURE analysis using PGDSpider v2.1.1.5 (Lischer & Excoffier, 2012). The number of putative clusters K was allowed to vary between 2 and 14, and chooseK.py was used to infer an optimal number of clusters maximising marginal likelihood.

## 3. RESULTS

We obtained 354,543,964 75-bp paired-end reads across 187 *Galaxias* specimens (Figure 1; Table S1). After filtering, we retained a dataset of 52,352 SNPs present in at least 90% of samples. Phylogenetic analyses of these data, with Australian *G. brevipinnis* samples specified as outgroups, recovered a well-resolved ML tree, with most major nodes receiving 100% bootstrap support (Figure 2, 3). The analysis provides support for the monophyly of NZ’s freshwater-limited *Galaxias vulgaris* complex (Figure 2, 3), with NZ *G. brevipinnis* (diadromous) placed sister to the stream-resident radiation. The analysis reveals substantial regional phylogenomic structure within the widespread *G. brevipinnis*, including individual monophyly of Tasmanian, mainland Australian, and NZ samples. The ‘trans-Tasman’ divergence between New Zealand and combined Australian *G. brevipinnis* samples is particularly strong (Figure 3). While no substantial phylogeographic structure is present among diadromous samples within these regions, fine-scale differentiation is evident for landlocked populations (e.g. subclades associated with different lakes: BT (Wakatipu), BO (Okareka), BZ (Great Lake)).

**FIGURE 3.**
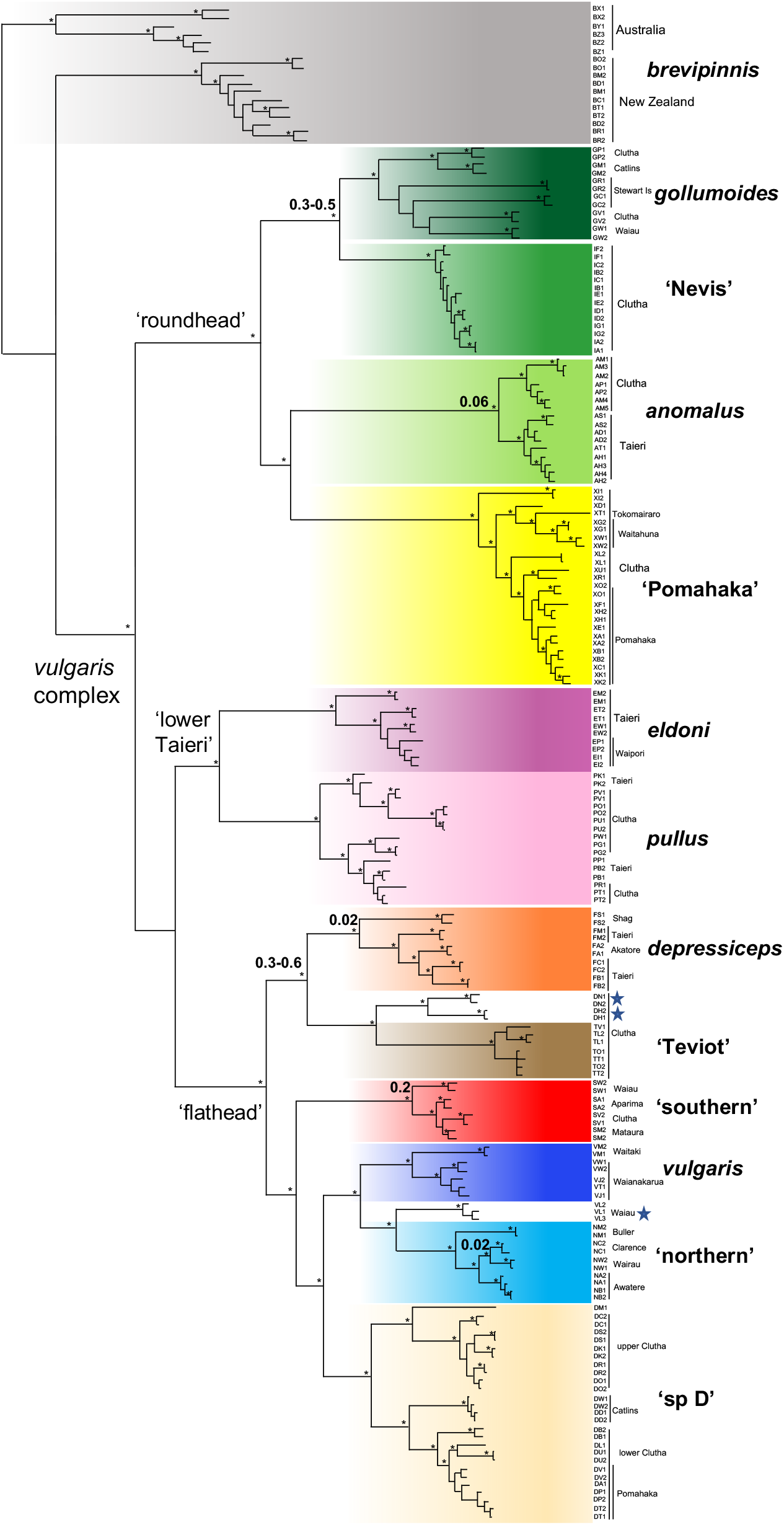
Maximum likelihood phylogeny of the stream resident *Galaxias vulgaris* complex, based on 52,352 SNPs, rooted with Australian samples of the diadromous sister species *Galaxias brevipinnis*. Nodes receiving >95% bootstrap support are indicated by small asterisks. Blue stars represent potential hybrid populations where lineages may have come into secondary contact (e.g. Esa et al., 2000). Estimated ages (mya) of divergence linked to geological river capture events are indicated in bold (see Waters et al., 2020).

Phylogenetic analyses based on both ML (Figure 3) and SNAPP (Figure S2) support the individual monophyly of all 11 previously recognized stream-resident lineages (typically with 100% bootstrap support; Figure 3), with a few very local exceptions. Specifically, four samples of *G*. ‘sp D’ (DH, DN) are placed sister to *G*. ‘Teviot’, likely reflecting hybridization among proximate ‘flathead’ lineages in secondary contact (Esa et al., 2000), and three northern samples of *G. vulgaris* (VL) are placed sister to the monophyletic *G*. ‘northern’. In addition to supporting the monophyly of currently recognized lineages, our analyses reveal a distinctive clade of stream-resident ‘roundhead’ samples centred on the Pomahaka R and adjacent regions of the lower Clutha system (which we here refer to as *G*. ‘Pomahaka’; Figures 1-3; Figure S2; see below). This clearly distinctive genomic lineage was previously unresolved by single-locus mtDNA analyses (Figure 2), as it carries mtDNA identical to that of the co-distributed ‘flathead’ *G*. ‘sp D’.

In terms of interspecific relationships, both maximum likelihood phylogenetic (Figure 3) and coalescent SNAPP (Figure S2) analyses of GBS loci support the monophyly of broad ‘flathead’, ‘roundhead’, and ‘lower Taieri’ (McDowall & Wallis, 1996; McDowall, 1997) species groups of stream-resident taxa. PCA (Figure 4) and fastSTRUCTURE (Figure S1) analyses similarly provide evidence for these broad species groupings, with particularly clear differentiation of ‘flathead’ and ‘roundhead’ species groups along PC1 (explanatory capacity 19.3%; Figure 4). The ‘flathead’ lineage cluster is resolved by fastSTRUCTURE at K = 3-6, and the ‘roundhead’ cluster at K = 2-3. Despite their relatively narrow geographic ranges (Figure 1), ‘roundhead’ lineages exhibit greater genetic divergence among one another than ‘flathead’ lineages (Figure 3, 4). This strong differentiation is further emphasised by the finding of multiple distinct ‘roundhead’ (but not ‘flathead’) clusters at higher values of K (Figure S1; optimal K =5).

**FIGURE 4.**
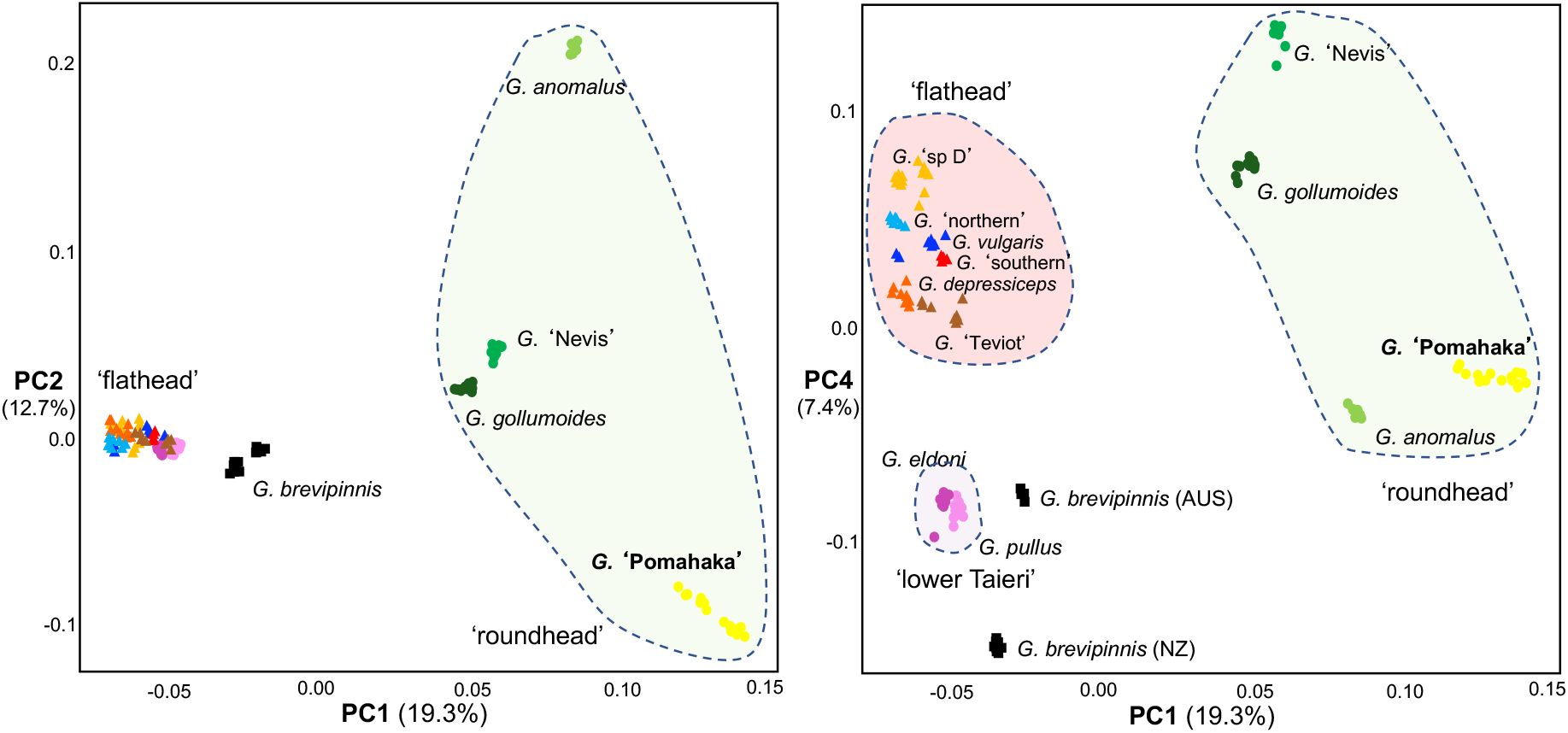
PCA plots based on 52,352 SNPs generated across 187 *Galaxias* specimens, showing relationships among 12 stream-resident lineages of the *G. vulgaris* complex (restricted to South Island, NZ) and diadromous *G. brevipinnis* (sampled from both NZ and Australia). Broad ‘flathead’ (triangles) versus ‘roundhead’ (circles) stream-resident species assemblages (indicated by large ellipses) are distinguishable on PC1. The newly-recognized *G*. ‘Pomahaka’ (‘roundhead’, lower Clutha system; shown in bold) was not resolved at the mtDNA level, but is clearly genomically distinct from the co-distributed and mitochondrially similar *G*. ‘sp D’ (Clutha ‘flathead’).

Phylogeographic analysis of genome-wide data highlights that the Clutha system has substantial *Galaxias* species diversity: eight of the 12 *vulgaris*-complex lineages (Figure 1). Furthermore, the Clutha system contains six tributary-specific lineages that are more closely related to lineages from neighbouring river systems. Specifically, the Nevis R (a Clutha tributary; Figure 1) houses a distinctive clade (*G*. ‘Nevis’; samples IA-IG) most closely related to Southland *G. gollumoides*, while the Teviot R (Figure 1) has a distinctive lineage (*G*. ‘Teviot’; TV, TT, TL, TO) that is sister to Taieri R *G. depressiceps* (Figure 1, 3). Similarly, the Von R (also a Clutha tributary) contains two species (GV, SV) otherwise broadly restricted to Southland rivers: *G. gollumoides* and *G*. ‘southern’ (Figure 1), while Clutha populations of *G. anomalus* (Manuherikia; AM, AP; Figure 1) and *G. pullus* (lower Clutha) have conspecific sister lineages in the Taieri R (Figure 3).

The lower Clutha system is dominated by two lineages: the ‘flathead’ lineage *G*. ‘sp D’ (throughout the Clutha system, including headwaters of the Pomahaka R: DP, DV, DT, DA), and *G*. ‘Pomahaka’, which is here resolved as a highly distinct ‘roundhead’ lineage (Figure 2, 3, 4; detected at K = 5 and K = 6; Figure S1) that is sister to *G. anomalus*, and completely unrelated to the co-distributed *G*. ‘sp D’ (despite their identical mtDNA). This newly-recognized ‘roundhead’ lineage is widespread across low-gradient streams of the Pomahaka R (Figure 1) and elsewhere in the lower Clutha system, with an additional isolated record from the adjacent Tokomairaro catchment (XT; Figure 1, 3). ABBA-BABA tests provide further evidence for extensive introgression between these co-distributed lineages (D statistic 0.27; *p* < 0.0001; Table S2).

Phylogeographic structuring within species occurs at a wide range of spatial scales, with multiple samples from single localities almost inevitably represented by monophyletic subclades (Figure 3). Such shallow structure occurs even among proximal locations within subcatchments (e.g. *G*. ‘Pomahaka’ samples XA, XB; *G*. ‘Nevis’ samples IA, IG, ID, IE, IF; *G. eldoni* samples EI, EP; *G. pullus* samples PO, PU; upper Clutha *G*. ‘sp D’ samples DC, DS, DK, DR, DO). Over broader spatial scales, the analysis frequently reveals substantial intraspecific phylogeographic structure *among* major sub-catchments and drainages. Such intraspecific phylogenomic ‘units’ include major subdivision detected within the widespread Clutha ‘flathead’ *G*. ‘sp D’, with distinct ‘upper Clutha’ (DR, DO, DC, DS, DK), ‘mid-Clutha (DM), ‘lower Clutha’ (DB, DU, DP, DV, DT, DA), and ‘Catlins’ (DW, DD) lineages (Figure 3). Strong river-specific and/or regional differentiation is also evident within *G. anomalus* (Taieri (AT, AH, AS, AD) versus Clutha (AM, AP)); for *G. depressiceps* (Taieri v Shag (FS)); *G. gollumoides* (Catlins (GW, GV) v Stewart Is (GR, GC) v Southland (GW, GV); and *G. vulgaris* (e.g. Waianakarua v Waitaki). By contrast, Taieri v Clutha populations of *G. pullus* are not reciprocally monophyletic.

## 4. DISCUSSION

### 4.1 Mitochondrial capture obscures cryptic diversity

Genome-wide analysis of the *G. vulgaris* complex provides support for 12 reciprocally-monophyletic freshwater-limited lineages in southern NZ, broadly consistent with previous phylogenetic analyses based on mtDNA (Waters & Wallis, 2001a, b) and nuclear markers (Waters et al., 2010; see Figure 2). However, one of these clades – *G*. ‘Pomahaka’ (lower Clutha roundhead) – was previously unrecognized, owing to the identical mtDNA lineage shared by it and the geographically co-distributed Clutha ‘flathead’ *G*. ‘sp D’ (Figure 2) (Burridge et al., 2007; Waters et al., 2001a; Waters et al., 2015b). While a preliminary nuclear DNA analysis alluded to a distinctive Pomahaka lineage (Waters et al., 2010), the current analysis highlights that *G*. ‘Pomahaka’ exhibits consistent genome-wide divergence relative to all other lineages and is widespread throughout the lower Clutha system (Figure 1, 3).

While there are several potential explanations for mito-nuclear discordance (e.g. selection; incomplete lineage sorting; introgression; Waters et al., 2010; Sunnucks et al., 2017; Wallis et al., 2017), introgression is a likely cause in the current system, given the extensive evidence for hybridization and introgression between lineages. In particular, the strong discordance between mtDNA versus genome-wide data for *G*. ‘Pomahaka’ (which carries mtDNA that is identical to that of the sympatric *G*. ‘sp D’, a species to which it is otherwise completely unrelated; Figure 2) is most simply explained by introgressive mitochondrial capture (see also Perea et al., 2016; Willis et al., 2014). Specifically, we infer that the native mtDNA of *G*. ‘Pomahaka’ has been replaced by mtDNA of sympatric *G*. ‘sp D’, due to hybridization (e.g. Perea et al., 2016), and this inference is further supported by ABBA-BABA tests for introgression between these co-distributed lineages (Table S2). Indeed, a growing number of freshwater phylogeographic studies have highlighted the remarkable potential for mtDNA introgression to generate extreme mito-nuclear discordance in such systems (Bisconti et al., 2018; Wallis et al., 2017). MtDNA can introgress among close sister taxa particularly readily, and while some such cases can be localised (e.g. Esa et al., 2000), others can lead to wholesale mtDNA replacement (Perea et al., 2016; Unmack et al., 2011; Willis et al., 2014). Consequently, such mtDNA capture has potential to substantially confound the accuracy of single-locus mtDNA barcoding analyses (approaches that have become increasingly popular in freshwater biodiversity studies; Bush et al., 2020; Hubert et al., 2008). In the current system, for instance, past reliance on mtDNA has apparently obscured the presence of a unique freshwater fish lineage, and such issues have potential to severely hamper conservation efforts. In a related example, diversity within the *G. olidus* complex was underestimated by mtDNA relative to nuclear allozymes, with introgressive hybridization and/or incomplete lineage sorting implicated (Adams et al., 2014). Moving forward, it is clear that the application of genome-wide molecular approaches is needed to resolve such mitonuclear discordance and facilitate meaningful detection and preservation of freshwater biodiversity (Buckley et al., 2018; Unmack et al., 2017).

Interestingly, our study also provides evidence of localized, asymmetric introgression of *G*. ‘sp D’ mtDNA into another ‘roundhead’ taxon, *G. anomalus*. This introgression is evident where these lineages co-occur in the Manuherikia R tributary of the Clutha (site AM/DM; Figure 1), and is supported also by ABBA-BABA analyses (Table S2). Specifically, two genomically *G. anomalus* specimens from this locality have yielded *G*. ‘sp D’ mtDNA (data not shown), whereas (as with *G*. ‘Pomahaka’) we have no evidence of roundhead mtDNA introgressing into *G*. ‘sp D’. We speculate that ecological preferences may render roundhead galaxiids particularly vulnerable to hybridization, with subsequent mtDNA introgression and capture, as their low-gradient habitats are more likely to experience disturbance, which can in turn promote hybridization (e.g. Grabenstein & Taylor, 2018). Such asymmetric mtDNA introgression may be enhanced by positive selection (e.g. Sunnucks et al., 2017) and/or genetic incompatibilities (Arntzen et al., 2009).

In addition to mtDNA replacement obscuring the diversity of taxa (above), our genome-wide analyses (Figure 2, 3) provide another example of putative mitochondrial capture influencing inferred relationships (see also MacGuigan & Near, 2019), with implications for our understanding of processes creating species diversity. Specifically, the mtDNA placement of *G. anomalus* as a close sister species of *G. pullus* (Figure 2; Waters & Wallis 2001a, b) strongly conflicts with both morphology (McDowall, 1997; McDowall & Wallis, 1996) and phylogenetic evidence from across the nuclear genome (Waters et al., 2010; current study). This mito-nuclear discordance likely stems from a relatively ancient introgressive mtDNA capture event, reflected by the approximately 2.0% divergence between control region of *G. anomalus* and *G. pullus*; Waters & Wallis, 2001b). Based on the phylogeographic history of these taxa (which currently co-occur in the Taieri R; Figure 1), we propose that historic introgressive replacement of *G. anomalus* mtDNA by *G. pullus* mtDNA occurred following the mid-late Pleistocene capture of the Kye Burn (formerly a tributary of the Clutha) by the proto-Taieri R (Craw et al., 2016; Waters et al., 2015a). Specifically, this composite formation of the Taieri likely mediated secondary contact and hybridization between *G. anomalus* (upper Taieri) and *G. pullus* (lower Taieri), apparently leading to introgressive mtDNA capture. Evidence for past introgression between these lineages is further provided by ABBA-BABA tests (Table S2). Hence, a mtDNA-centric assessment would invoke diversification of these taxa within the Taieri, as opposed to the invasion of one species from a neighbouring catchment.

### 4.2 Population translocation, fragmentation and conservation

Human impacts have potential to erode and reshape freshwater biodiversity. As a case in point, artificial translocation events can promote hybridization and introgression between formerly isolated lineages (e.g. Blackwell et al., 2021; DeMarais et al., 1992; Echelle & Echelle, 1997), with potential implications for conservation (McDowall 1990). Our genetic analyses have highlighted the potential impacts of man-made drainage alterations (e.g. water races, impoundments) on fish distributions in NZ (Esa et al., 2000) and elsewhere (Waters et al., 2002). In NZ, gold-mining operations in the 1800s often relied on construction of artificial water race connections between the headwaters of adjacent drainage systems (Esa et al., 2000), potentially connecting disjunct freshwater lineages. The hypothesis of anthropogenic secondary contact as a mediator of galaxiid hybridization is supported in the current study by the localised finding of an apparent hybrid genotype in the historic Poolburn water race (DN) which artificially connects adjacent Taieri (*G. depressiceps*) and Manuherikia (*G*. ‘sp D’) ‘flathead’ populations otherwise separated by Rough Ridge (Figure 1, 3; Esa et al., 2000). The current study provides additional evidence for similar anthropogenic translocation events elsewhere, including the distribution of a *G*. ‘Pomahaka’ subclade (XT, XG, XW; Figure 1, 3) across the low Clutha-Tokomairaro divide (where an historic water race connected the adjacent Waitahuna (Clutha system; XG, XW) and Manuka (Tokomairaro R; XT) streams). Indeed, the Manuka Stream record represents the only known occurrence of *G*. ‘Pomahaka’ outside the Clutha system, and this local translocation scenario is supported by sub-catchment phylogeographic structure within this lineage (Figure 3). Comparable artificial water race connections seem likely to also explain the genetic similarity *G. pullus* populations from adjacent headwaters of the Tuapeka (Clutha; PT, PR) and Waipori (Taieri; PB, PP) systems.

In addition to anthropogenic translocations and hybridization, NZ’s stream-resident galaxiids are also threatened by population fragmentation, which has arisen via a combination of habitat modification (e.g. through impoundments) and invasive species introductions over the last 150 years (e.g. trout; McDowall, 1990, 2006). Such lasting fragmentation may lead to rapid genetic drift and even population extirpation. Notably, our genome-wide study detected strong phylogeographic structure within species, even among adjacent sites within single sub-catchments (e.g. *G*. ‘Nevis’ samples IA, IG, ID, IE, IF; *G. eldoni* samples EI, EP). While the relatively deep phylogeographic structures we observe over broad spatial scales clearly have pre-human origins (e.g. based on deep mtDNA and genome-wide divergence between upper versus lower Clutha clades of *G*. ‘sp D’; Burridge et al., 2007; Waters et al., 2001a), we suspect that many shallow divergences (and reciprocal monophyly) detected among neighbouring sites within rivers at least partly reflect anthropogenic population fragmentation. For instance, reciprocally monophyletic samples of upper Clutha *G*. ‘sp D’ (DR, DO, DC, DS, DK) are isolated by both an impoundment and by introduced salmonids; the situation is similar for Waipori R populations of *G. eldoni* (EI, EP) (Figure 1, 3). Broadly, a combination of habitat fragmentation and predation by introduced salmonids (McDowall, 1990, 2006) has resulted in the widespread decline and/or extirpation of numerous *G. vulgaris* complex populations, which are now broadly restricted to isolated, trout-free habitats (Townsend & Crowl, 1991). This recent fragmentation, and lack of main-channel population connectivity, is highlighted here by the evolution of shallow genome-wide phylogeographic structure among proximate populations that were presumably connected prior to trout introduction. Given the isolating role of invasive salmonids (McDowall, 2006; Townsend & Crowl, 1991), there is an increasing need to locally remove trout, and to install trout barriers, to allow for recovery and re-connectivity of native galaxiid populations (e.g. Lintermans, 2000). Additionally, human-mediated galaxiid gene flow should be considered in areas where anthropogenically isolated populations are potentially experiencing inbreeding depression (i.e. ‘genetic rescue’; Burridge, 2019; Whiteley et al., 2015).

### 4.3 River capture and fish evolution

Geological processes affecting river drainage connectivity are thought to have played important roles in the divergence of freshwater-limited lineages in many regions of the globe (e.g. Craw et al., 2015; Goodier et al., 2011; Kozak et al., 2006; Mayden, 1988). In several cases, the persistent genetic legacies of palaeodrainage features have been highlighted by interdisciplinary studies combining geological and biological data (e.g. Waters et al., 2020b). In the current study, we show that the Clutha River is home to a particularly diverse suite of *Galaxias vulgaris* complex clades (including eight of 12 currently recognized stream resident lineages; Figure 1), likely reflecting the river’s complex, composite geological formation (Craw et al., 2012). Indeed, our galaxiid GBS analyses provide genome-wide support for several previously-proposed links between fish biogeography and river capture. Specifically, several headwater tributaries of the Clutha River system harbour distinctive *Galaxias* lineages apparently ‘captured’ geologically from neighbouring drainage systems. In particular, genome-wide data support geologically-proposed paleodrainage connections between the Nevis (*G*. ‘Nevis’) and Mataura (Southland, *G. gollumoides*) (Waters et al., 2001a); between the Teviot (*G*. ‘Teviot’) and Taieri (*G. depressiceps*) (Waters et al., 2015a); between the Von and Oreti (related populations of *G. gollumoides* and *G*. ‘southern’; Burridge et al., 2007); and between the Manuherikia and Taieri (divergent populations of *G. anomalus*; Craw et al., 2007).

The current study similarly finds support for dynamic river geological histories elsewhere in NZ. Notably, genome-wide analyses support a geologically composite origin for Otago’s Taieri R (Craw et al., 2016; Waters et al., 2015), evidenced by divergent ‘upper Taieri’ (*G. anomalus, G. depressiceps*) versus ‘lower Taieri’ (*G. eldoni, G. pullus*; Figures 1-3) fish assemblages. Likewise, our genetic data similarly support sister relationships between Taieri and Shag R *G. depressiceps* (Craw et al., 2016), a geologically-derived capture hypothesis previously supported by mtDNA alone. Finally, the sister relationship observed for Wairau (NW) and Clarence (NC) populations of *G*. ‘northern’, relative to samples from the geographically intermediate Awatere R (NA, NB), strongly supports the proposed glacially-mediated capture event involving the headwaters of the two former rivers (Burridge et al., 2006; McAlpin, 1992).

### 4.4 Ecological and phylogeographic diversification

The phylogenetic detection of ecologically and morphologically distinct ‘roundhead’ (limnetic; pool-dwelling; surface feeding) versus ‘flathead’ (benthic; riffle-dwelling, midwater feeding) species groups (Figures 3, 4) suggests that the broad divergence of these ecomorphs (e.g. Crow et al., 2009; Crow 2010) was initiated relatively early in the radiation of the *G. vulgaris* complex. Additionally, the finding that genetic distances among ‘roundhead’ lineages exceed those detected among ‘flathead’ lineages (Figure 3, 4) may partly reflect the relatively deep phylogenetic divergence of the former (despite their relatively narrow distributions). We also speculate that ecological preferences for slow-flowing pools and backwaters may increase rates of evolutionary divergence among ‘roundhead’ lineages (i.e. population fragmentation due to the absence of main-channel connectivity; Waters & Burridge, 2016). While the ‘roundhead’ clade has not achieved the dramatic northward biogeographic expansion evident in the more-recent ‘flathead’ radiation (Figure 1, 3), it nevertheless shows evidence of localised movement across low drainage divides (Burridge et al., 2008b).

### 4.5 Conclusions

In summary, the current study highlights the power of high-throughput sequencing approaches to resolve the phylogenies of species-rich freshwater radiations, and for addressing long-standing systematic puzzles arising from mitonuclear discordance. Our study provides a key example of mtDNA capture which illustrates the need for multi-locus approaches to accurately characterize freshwater biodiversity. Based on these data we also infer the role of natural and anthropogenic drainage alterations involved in the dispersal, mixing and diversification of freshwater-limited lineages (Allibone & Wallis, 1993; Waters et al., 2020a). Such hybridization and translocation events present potential threats to species conservation (Rhymer & Simberloff, 1996). Overall, it is clear that genome-wide approaches are crucial for resolving and conserving freshwater biodiversity.

## Acknowledgments

We wish to acknowledge the use of NZ eScience Infrastructure (NeSI) high performance computing facilities, consulting support and training services as part of this research. NZ’s national facilities are provided by NeSI and funded jointly by NeSI’s collaborator institutions and through the Ministry of Business, Innovation & Employment’s Research Infrastructure programme. Florian Privé provided very helpful analytical input. We also acknowledge crucial support from the Marsden fund and NZ Department of Conservation.

---

**Table S1.**
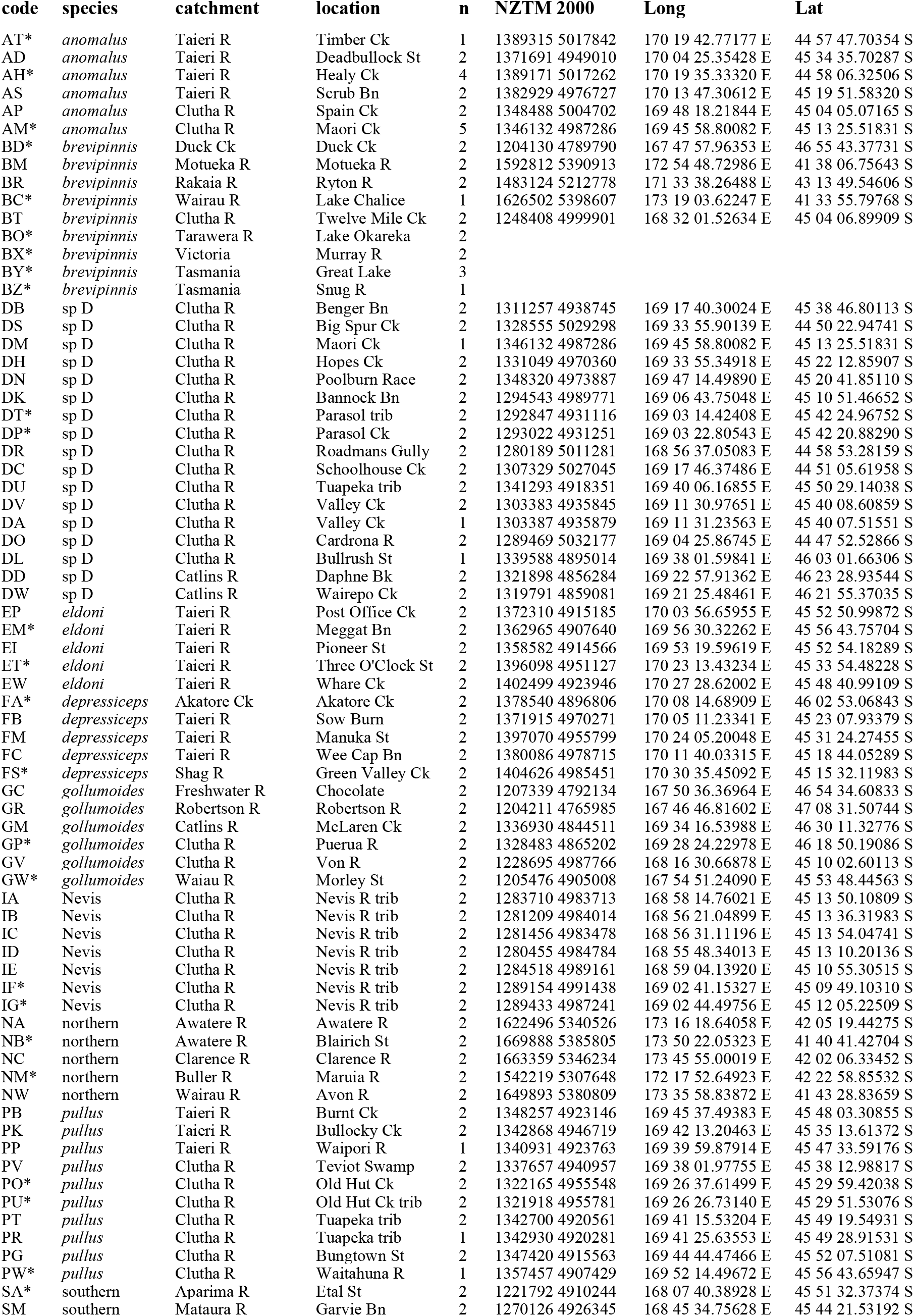
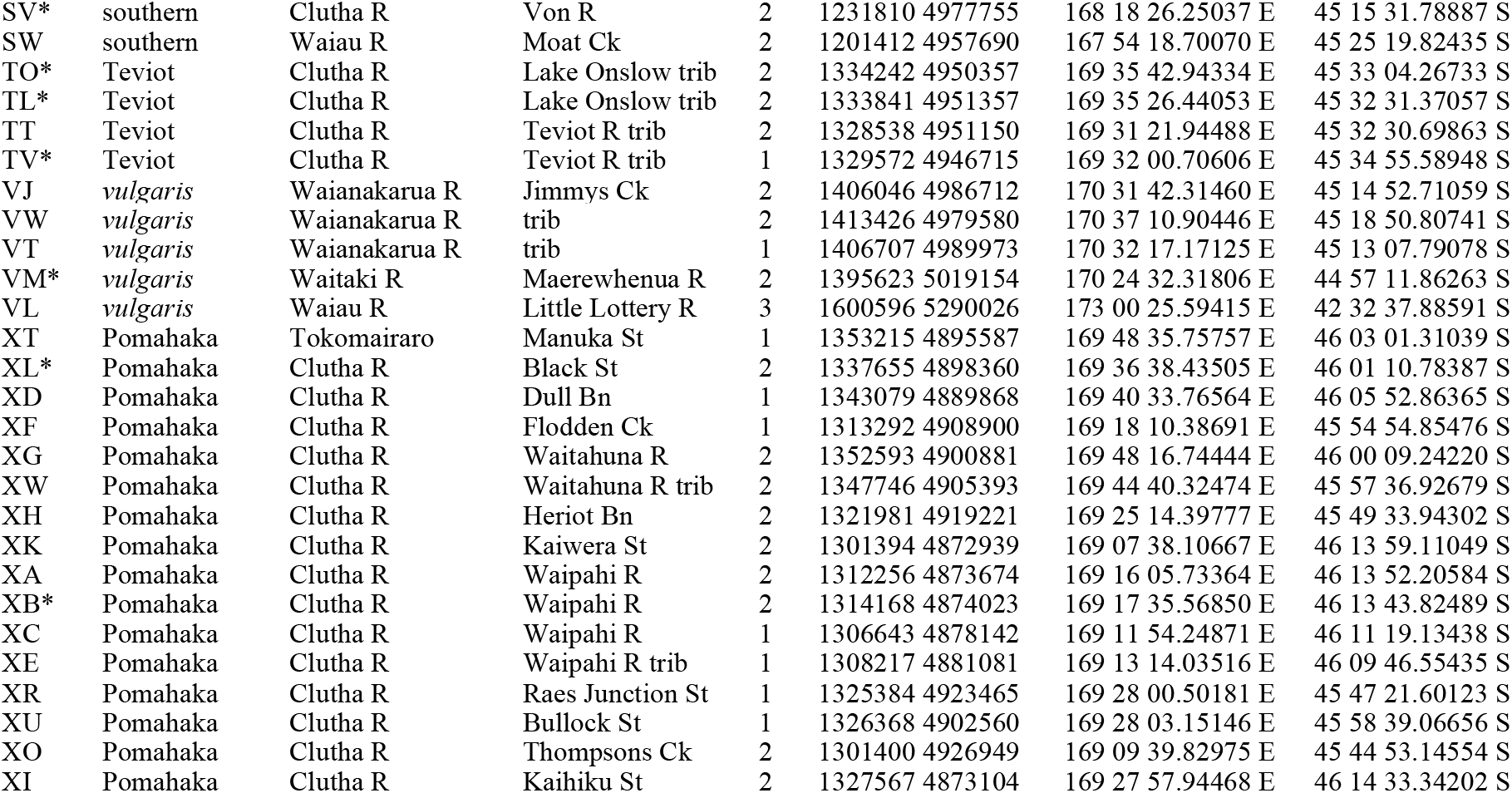
Collection details for *Galaxias* specimens included in genome-wide analyses. Asterisks indicate samples included in SNAPP analyses (Figure S2).

**Table S2.** ABBA-BABA statistics for selected stream-resident *Galaxias* lineages, using New Zealand *Galaxias brevipinnis* as an outgroup. An excess of ABBA over BABA indicates introgression between lineages two and three, whereas an excess of BABA over ABBA would indicate introgression between lineages one and three. All three tests reveal highly significant introgression between lineages 2 and 3.

| Lineage 1 | Lineage 2 | Lineage 3 | D-statistic | p-value | BBAA | ABBA | BABA |
| --- | --- | --- | --- | --- | --- | --- | --- |
| *G. gollumoides* | *G. 'Pomahaka'* | *G. 'sp D'* | 0.27 | $< 6.68 \times 10^{-8}$ | 823.67 | 198.57 | 115.27 |
| *G. gollumoides* | *G. anomalus* | *G. 'sp D'* | 0.36 | $< 2.30 \times 10^{-16}$ | 726.34 | 266.37 | 124.75 |
| *G. gollumoides* | *G. anomalus* | *G. pullus* | 0.18 | $2.89 \times 10^{-4}$ | 955.05 | 146.60 | 101.84 |

**FIGURE S1.**
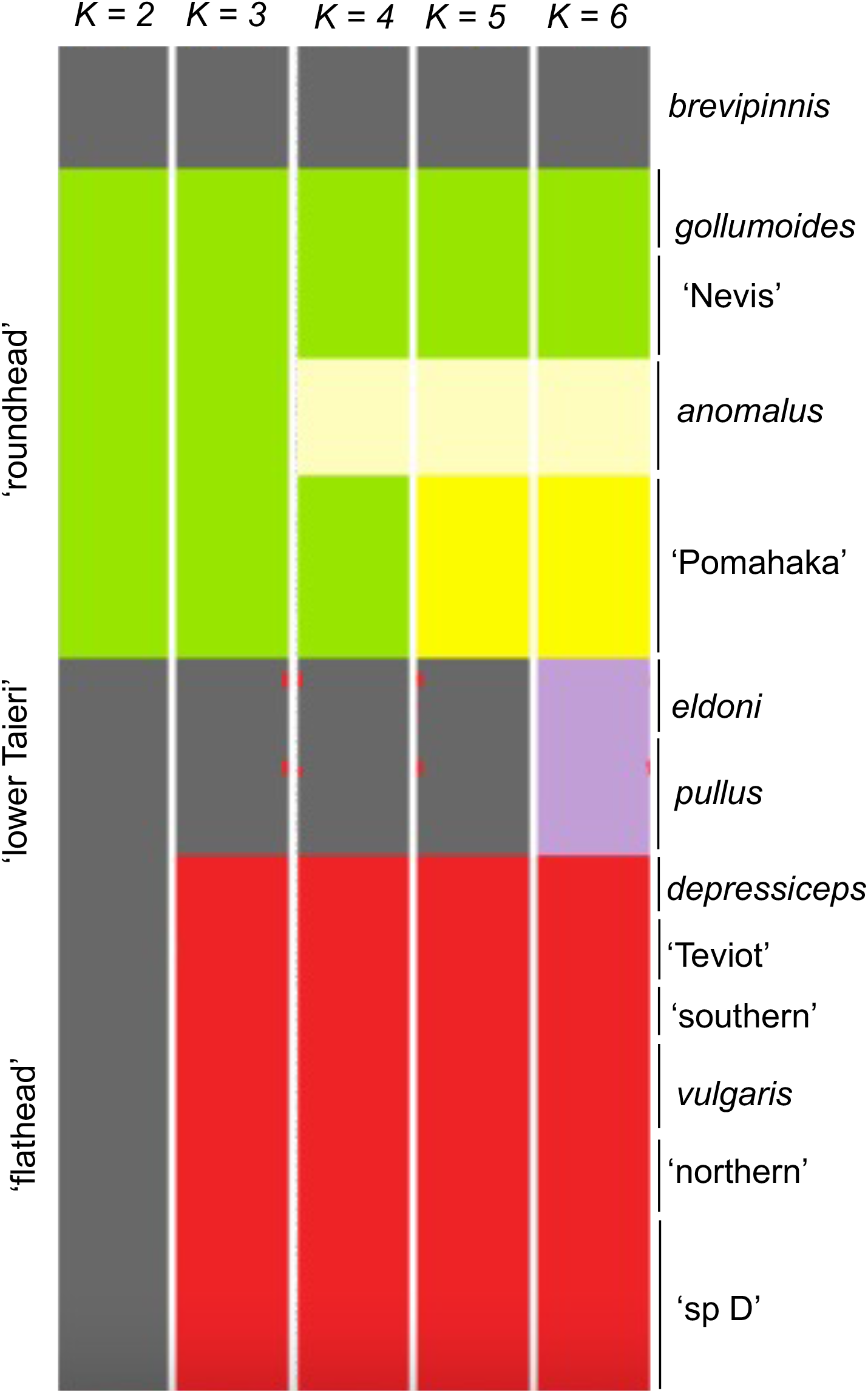
FastSTRUCTURE analyses of the *Galaxias vulgaris* complex based on genome-wide SNP data, with marginal likelihood maximized at K = 5. Major species groupings are resolved (labels on left), and the newly recognized ‘roundhead’ lineage *Galaxias* ‘Pomahaka’ (yellow) is supported as a separate genomic cluster at K = 5 and K = 6, being clearly distinct from the co-distributed ‘flathead’ lineage *Galaxias* ‘sp D’ (red).

**FIGURE S2.**
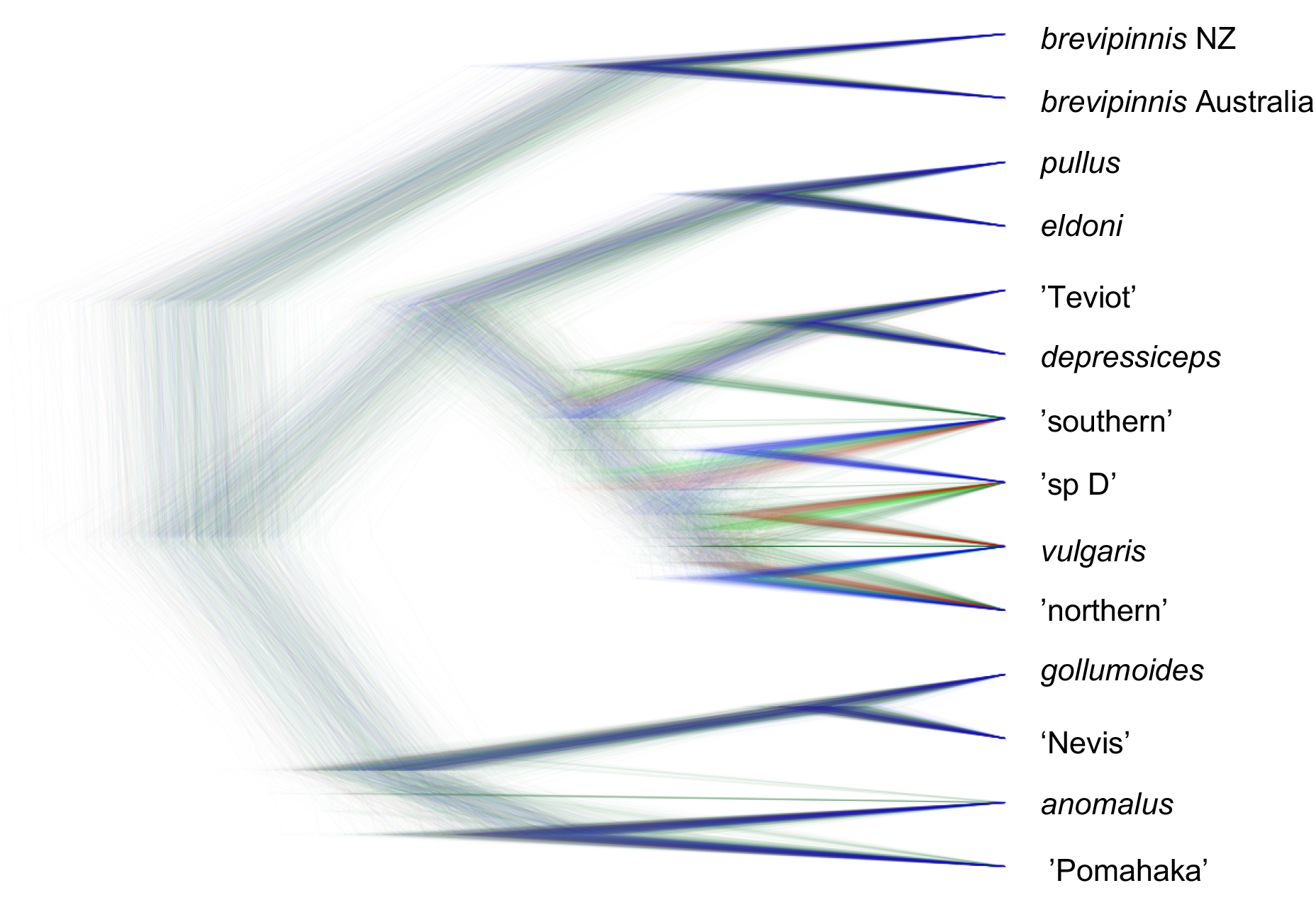
Coalescent phylogenetic analysis of stream resident *Galaxias* based on unlinked SNPs, performed using SNAPP (representing 2000 trees). The analysis was rooted with diadromous *Galaxias brevipinnis* as an outgroup. The most common topology is represented in blue, followed by the next most common trees in red and then dark green. All the other trees are in light green.

## REFERENCES

Adams, M., Raadik, T. A., Burridge, C. P., & Georges, A. (2014). Global biodiversity assessment and hyper-cryptic species complexes: more than one species of elephant in the room? Systematic Biology, 63, 518–33.

Allendorf, F. W., Funk, W. C., Aitken, S. N., Byrne, M., & Luikart, G. (2022). Conservation and the Genomics of Populations. 3rd Edition, Oxford University Press.

Allibone, R. M., Crowl, T. A., Holmes, J. M., King, T. M., McDowall, R. M., Townsend, C. R., & Wallis, G. P. (1996). Isozyme analysis of Galaxias species (Teleostei: Galaxiidae) from the Taieri River, South Island, New Zealand: a species complex revealed. Biological Journal of the Linnean Society, 57, 107–127.

Allibone, R. M., & Townsend, C. R. (1997). The distribution of four recently discovered galaxiid species in the Taieri River, New Zealand: the role of microhabitat. Journal of Fish Biology, 51, 1235–1246.

Allibone, R. M., & Wallis, G. P. (1993). Genetic variation and diadromy in some native New Zealand galaxiids (Teleostei: Galaxiidae). Biological Journal of the Linnean Society, 50, 19–33.

Andrews, K. R., Good, J. M., Miller, M. R., Luikart, G., & Hohenlohe, P. A. (2016). Harnessing the power of RADseq for ecological and evolutionary genomics. Nature Reviews Genetics, 17, 81–92.

Arntzen, J. W., Jehle, R., Bardacki, F., Burke, T., & Wallis, G. P. (2009). Asymmetric viability of reciprocal-cross hybrids between crested and marbled newts (Triturus cristatus and T. marmoratus). Evolution, 63, 1191–1202.

Baird, N. A., Etter, P. D., Atwood, T. S., Currey, M. C., Shiver, A. L., & Lewis, Z. A. (2008) Rapid SNP discovery and genetic mapping using sequenced RAD markers. PLoS ONE, 3, e3376.

Barluenga, M., Stolting, K. N., Salzburger, W., Muschick, M., & Meyer, A. (2006). Sympatric speciation in Nicaraguan crater lake cichlid fish. Nature, 439, 719–723.

Bisconti, R., Porretta, D., Arduino, P., Nascetti, G., & Canestrelli, D. (2018). Hybridization and extensive mitochondrial introgression among fire salamanders in peninsular Italy. Scientific Reports, 8, 13187.

Blackwell, T., Ford, A. G., Ciezarek, A. G., Bradbeer, S. J., Gracida Juarez, C. A., Smith, A. M., Ngatunga, B. P., Shechonge, A., Tamatamah, R., & Etherington, G. (2021). Newly discovered cichlid fish biodiversity threatened by hybridization with non-native species. Molecular Ecology, 30, 895–911.

Bouckaert, R. (2010). DensiTree: making sense of sets of phylogenetic trees. Bioinformatics, 26, 1372–1373.

Bouckaert R., Vaughan, T. G., Barido-Sottani, J., Duchêne, S., Fourment, M., Gavryushkina A., et al. (2019). BEAST 2.5: An advanced software platform for Bayesian evolutionary analysis. PLoS Computational Biology, 15, e1006650.

Bryant, D., Bouckaert, R., Felsenstein, J., Rosenberg, N. A., & RoyChoudhury, A. (2012). Inferring species trees directly from biallelic genetic markers: Bypassing gene trees in a full coalescent analysis. Molecular Biology and Evolution, 29, 1917–1932.

Buckley, S. J., Domingos, F., Attard, C. R. M., Brauer, C. J., Sandoval-Castillo, J., Lodge, R., Unmack, P. J., & Beheregaray, L. B. (2018). Phylogenomic history of enigmatic pygmy perches: implications for biogeography, taxonomy and conservation. Royal Society Open Science, 5, 172125.

Burridge, C. P. (2019). Politics and pride: maintaining genetic novelty may be detrimental for the conservation of Formosa landlocked salmon Oncorhynchus formosanus. Aquatic Conservation: Marine and Freshwater Ecosystems, 29, 840–847.

Burridge, C. P., Craw, D., Fletcher, D., & Waters, J. M. (2008a). Geological dates and molecular rates: fish DNA sheds light on time dependency. Molecular Biology and Evolution, 25, 624–33.

Burridge, C. P., Craw, D., & Waters, J. M. (2007). An empirical test of freshwater vicariance via river capture. Molecular Ecology, 16, 1883–1895.

Burridge, C. P., Craw, D., Jack, D. C., King, T. M., & Waters, J. M. (2008b). Does fish ecology predict dispersal across a river drainage divide? Evolution, 62, 1484–1499.

Burridge, C. P., Craw, D., & Waters, J. M. (2006). River capture, range expansion, and cladogenesis: the genetic signature of freshwater vicariance. Evolution, 60, 1038–1049.

Burridge, C. P., McDowall, R. M., Craw, D., Wilson, M. V. H., & Waters, J. M. (2012). Marine dispersal as a pre-requisite for Gondwanan vicariance among elements of the galaxiid fish fauna. Journal of Biogeography, 39, 306–321.

Burridge, C. P., & Waters, J. M. (2020). Does migration promote or inhibit diversification? A case study involving the dominant radiation of temperate Southern Hemisphere freshwater fishes. Evolution, 74, 1954–1965.

Bush, A., Monk, W. A., Compson, Z. G., Peters, D. L., Porter, T. M., Shokralla, S., Wright, M. T. G., Hajibabaei, M., & Baird, D. J. (2020). DNA metabarcoding reveals metacommunity dynamics in a threatened boreal wetland wilderness. Proceedings of the National Academy of Sciences USA, 117, 8539–8545.

Catchen, J., Hohenlohe, P. A., Bassham, S., Amores, A., & Cresko, W. A. (2013). Stacks: an analysis tool set for population genomics. Molecular Ecology, 22, 3124–3140.

Chakona, A., Swartz, E. R., & Gouws, G. (2013). Evolutionary drivers of diversification and distribution of a southern temperate stream fish assemblage: testing the role of historical isolation and spatial range expansion. PLoS One, 8, e70953.

Closs, G. P., Krkosek, M., & Olden, J. D. (2016). Conservation of Freshwater Fishes. Cambridge University Press.

Colosimo, P. F., Hosemann, K. E., Balabhadra, S., Villarreal, G., Dickson, M., Grimwood, J., Schmutz, J., Myers, R. M., Schluter, D., & Kingsley, D. M. (2005). Widespread parallel evolution in sticklebacks by repeated fixation of ectodysplasin alleles. Science, 307, 1928–1933.

Craw, D., Burridge, C. P., & Waters, J. M. (2007). Geological and biological evidence for drainage reorientation during uplift of alluvial basins, central Otago, New Zealand. New Zealand Journal of Geology and Geophysics, 50, 367–376.

Craw, D., Craw, L., Burridge, C. P., Wallis, G. P., & Waters, J. M. (2016). Evolution of the Taieri River catchment, East Otago, New Zealand. New Zealand Journal of Geology and Geophysics, 59, 257–273.

Craw, D., Upton, P., Burridge, C. P., Wallis, G. P., & Waters, J. M. (2015). Rapid biological speciation driven by tectonic evolution in New Zealand. Nature Geoscience, 9, 140–144.

Craw, D., Upton, P., Walcott, R., Burridge, C. P., & Waters, J. M. (2012). Tectonic controls on the evolution of the Clutha River catchment, New Zealand. New Zealand Journal of Geology and Geophysics, 55, 345–359.

Crow, S. K., Closs, G. P. Waters, J. M., Booker, D. J., & Wallis, G. P. (2010). Niche partitioning and the effect of interspecific competition on microhabitat use by two sympatric galaxiid stream fishes. Freshwater Biology, 55, 967–982.

Crow, S. K., Waters, J. M., Closs, G. P., & Wallis, G. P. (2009). Morphological and genetic analysis of Galaxias ‘southern’ and G. gollumoides: interspecific differentiation and intraspecific structuring. Journal of the Royal Society of New Zealand, 39, 43–62.

Danecek, P., Auton, A., Abecasis, G., Albers, C. A., Banks, E., Depristo, M. A., Handsaker, R. E., Lunter, G., Marth, G. T., & Sherry, S. T. (2011). The variant call format and VCFtools. Bioinformatics, 27, 2156–2158.

Delgado, M. L., Górski, K., Habit, E., & Ruzzante, D. E. (2019). The effects of diadromy and its loss on genomic divergence: the case of amphidromous Galaxias maculatus populations. Molecular Ecology, 28, 5217–5231.

Delgado, M. L., Manosalva, A., Urbina, M. A., Habit, E., Link, O., & Ruzzante, D. E. (2020). Genomic basis of the loss of diadromy in Galaxias maculatus: insights from reciprocal transplant experiments. Molecular Ecology, 29, 4857–4870.

DeMarais, B. D., Dowling, T. E., Douglas, M. E., Minckley, W. L., & Marsh, P. C. (1992). Origin of Gila seminuda (Teleostei: Cyprinidae) through introgressive hybridization: implications for evolution and conservation. Proceedings of the National Academy of Sciences USA, 117, 89, 2747–2751.

Dray, S., & Dufour, A. (2007). The ade4 package: implementing the duality diagram for ecologists. Journal of Statistical Software, 22, 1–20.

Dunn, N. R., Allibone, R. M., Closs, G., Crow, S., David, B. O., Goodman, J., Griffiths, M. H., Jack, D. C., Ling, N., & Waters, J. M. (2018). Conservation status of New Zealand freshwater fishes, 2017. NZ Department of Conservation.

Durand, E. Y., Patterson, N., Reich, D., & Slatkin, M. (2011). Testing for ancient admixture between closely related populations. Molecular Biology and Evolution, 28, 2239–2252.

Echelle, A. A., & Echelle, A. F. (1997). Genetic introgression of endemic taxa by non-natives: a case study with Leon Springs pupfish and sheepshead minnow. Conservation Biology, 11, 153–161.

Elshire, R. J., Glaubitz, J. C., Sun, Q., Poland, J. A., Kawamoto, K., Buckler, E. S. & Mitchell, S. E. (2011). A robust, simple genotyping-by-sequencing (GBS) approach for high diversity species. PLoS One, 6, e19379.

Esa, Y. B., Waters, J. M., & Wallis, G. P. 2000. Introgressive hybridization between Galaxias depressiceps and Galaxias sp D (Teleostei: Galaxiidae) in Otago, New Zealand: secondary contact mediated by water races. Conservation Genetics, 1, 329–339.

Fang, B., Kemppainen, P., Momigliano, P., Feng, X., & Merilä, J. (2020). On the causes of geographically heterogeneous parallel evolution in sticklebacks. Nature Ecology and Evolution, 4, 1105–1115.

Ford, A. G. P., Bullen, T. R., Pang, L., Genner, M. J., Bills, R., Flouri, T., Ngatunga, B. P., Rüber, L., Schliewen, U. K., Seehausen, O., Shechonge, A., Stiassny, M. L. J., Turner, G. F., & Day, J. J. (2019). Molecular phylogeny of Oreochromis (Cichlidae: Oreochromini) reveals mito-nuclear discordance and multiple colonisation of adverse aquatic environments. Molecular Phylogenetics and Evolution, 136, 215–226.

Goodier, S. A. M., Cotterill, F. P. D., O’Ryan, C., Skelton, P. H., & de Wit, M. J. (2011). Cryptic diversity of African tigerfish (genus Hydrocynus) reveals palaeogeographic signatures of linked Neogene geotectonic events. PLoS One, 6, e28775.

Grabenstein, K. C., & Taylor, S. A. (2018). Breaking barriers: causes, consequences, and experimental utility of human-mediated hybridization. Trends in Ecology and Evolution, 33, 198–212.

Graham, C. F., Glenn, T. C., McArthur, A. G., Boreham, D. R., Kieran, T., Lance, S., Manzon, R. G., Martino, J. A., Pierson, T., & Rogers, S. M. (2015). Impacts of degraded DNA on restriction enzyme associated DNA sequencing (RADSeq). Molecular Ecology Resources, 15, 1304–1315.

Green, R. E., Krause, J., Briggs, A. W., Maricic, T., Stenzel, U., Kircher, M., Patterson, N., Li, H., Zhai, W., Fritz, M. H. Y., Hansen, N. F., Durand, E. Y., Malaspinas, A. S., Jensen, J. D., Marques-Bonet, T., Alkan, C., Prufer, K., Meyer, M., Burbano, H. A., … Paabo, S. (2010). A draft sequence of the Neandertal genome. Science, 328, 710–722.

Hubert, N., Hanner, R., Holm, E., Mandrak, N. E., Taylor, E., Burridge, M., Watkinson, D., Dumont, P., Curry, A., Bentzen, P., Zhang, J., April, J., & Bernatchez, L. (2008). Identifying Canadian freshwater fishes through DNA barcodes. PLoS One, 3, e2490.

Irisarri, I., Singh, P., Koblmüller, S. et al. (2018). Phylogenomics uncovers early hybridization and adaptive loci shaping the radiation of Lake Tanganyika cichlid fishes. Nature Communications, 9, 3159.

Jombart, T. (2008). adegenet: a R package for the multivariate analysis of genetic markers. Bioinformatics, 24, 1403–1405.

Jones, P. E., & Closs, G. P. (2016). Interspecific differences in early life-history traits in a species complex of stream-resident galaxiids. Ecology of Freshwater Fish, 25, 211–224.

Jones, P. E., Senior, A., Allibone, R. M., & Closs, G. P. (2016). Life history variation in a species complex of nonmigratory galaxiids. Ecology of Freshwater Fish, 25, 174–189.

Kamvar, Z. N., Tabima, J. F., & Grünwald, N. J. (2014). Poppr: an R package for genetic analysis of populations with clonal, partially clonal, and/or sexual reproduction. PeerJ, 2, e281.

Knaus, B. J., & Grünwald, N. J. (2017). vcfr: a package to manipulate and visualize variant call format data in R. Molecular Ecology Resources, 17, 44–53.

Kozak, K. H., Blaine, R. A., & Larson, A. (2006). Gene lineages and eastern North American palaeodrainage basins: phylogeography and speciation in salamanders of the Eurycea bislineata species complex. Molecular Ecology, 15, 191–207.

Lintermans, M. (2000). Recolonization by the mountain galaxias Galaxias olidus of a montane stream after the eradication of rainbow trout Oncorhynchus mykiss. Marine and Freshwater Research, 51, 799–804.

Lischer, H. E. L., & Excoffier, L. (2012). PGDSpider: an automated data conversion tool for connecting population genetics and genomics programs. Bioinformatics, 28, 298–299.

Luu, K., Bazin, E., & Blum, M. G. B. (2017). pcadapt: an R package to perform genome scans for selection based on principal component analysis. Molecular Ecology Resources, 17, 67–77.

Malinsky, M., Matschiner, M., & Svardal, H. (2021). Dsuite - Fast D-statistics and related admixture evidence from VCF files. Molecular Ecology Resources, 21, 584–595.

Mayden, R. L. (1988). Vicariance biogeography, parsimony, and evolution in North American freshwater fishes. Systematic Zoology, 37, 329–355.

MacGuigan, D. J., & Near, T. J. (2019). Phylogenomic signatures of ancient introgression in a rogue lineage of darters (Teleostei: Percidae). Systematic Biology, 68, 329–346.

McAlpin, J. P. (1992). Glacial geology of the upper Wairau Valley, Marlborough, New Zealand. New Zealand Journal of Geology and Geophysics, 35, 211–222.

McDowall, R. M. (1970). The galaxiid fishes of New Zealand. Bulletin of the Museum of Comparative Zoology, 139, 341–431.

McDowall, R. M. (1990). New Zealand Freshwater Fishes: a Natural History and Guide. Heinemann Reid, MAF Publishing Group.

McDowall, R. M. (2006). Crying wolf, crying foul, or crying shame: alien salmonids and a biodiversity crisis in the southern cool-temperate galaxioid fishes? Reviews in Fish Biology and Fisheries, 16, 233–422.

McDowall, R. M. (1997). Two further new species of Galaxias (Teleostei: Galaxiidae) from the Taieri River, southern New Zealand. Journal of the Royal Society of New Zealand, 27, 199–217.

McDowall, R. M., & Wallis, G. P. (1996). Description and redescription of Galaxias species (Teleostei: Galaxiidae) from Otago and Southland. Journal of the Royal Society of New Zealand, 26, 401–427.

Melo, B. F., Sidlauskas, B. L., Near, T. J., Roxo, F. F., Ghezelayagh, A., Ochoa, L. E., Stiassny, M. L. J., Arroyave, J., Chang, J., Faircloth, B. C., MacGuigan, D. J., Harrington, R. C., Benine, R. C., Burns, M. D., Hoekzema, K., Sanches, N. C., Maldonado-Ocampo, J. A., Castro, R. M. C., Foresti, F., Alfaro, M. E., & Oliveira, C. (2022). Accelerated diversification explains the exceptional species richness of tropical characoid fishes. Systematic Biology, 71, 78–92.

Olden, J. D., Kennard, M. J., Leprieur, F., Tedesco, P. A., Winemiller, K. O., & Garcia-Bethou, E. (2010). Conservation biogeography of freshwater fishes: recent progress and future challenges. Diversity and Distributions, 16, 496–513.

Perea, S., Vukić, J., Šanda, R., & Doadrio, I. (2016). Ancient mitochondrial capture as factor promoting mitonuclear discordance in freshwater fishes: a case study in the genus Squalius (Actinopterygii, Cyprinidae) in Greece. PLoS One, 11, e0166292.

R Core Team. (2020). R: a language and environment for statistical computing. R Foundation for Statistical Computing website.

Raadik, T. A. (2014). Fifteen from one: a revision of the Galaxias olidus Gunther, 1866 complex (Teleostei, Galaxiidae) in south-eastern Australia recognises three previously described taxa and describes 12 new species. Zootaxa, 3898, 1–198.

Raj, A., Stephens, M., & Pritchard, J. K. (2014). fastSTRUCTURE: variational inference of population structure in large SNP data sets. Genetics, 197, 573–589.

Rambaut, A. (2009). FigTree. Tree figure drawing tool. http://tree.bio.ed.ac.uk/software/figtree/

Rhymer, J. M., & Simberloff, D. (1996). Extinction by hybridization and introgression. Annual Review of Ecology and Systematics, 27, 83–109.

Ronco, F., Matschiner, M., Böhne, A. et al. (2021) Drivers and dynamics of a massive adaptive radiation in cichlid fishes. Nature, 589, 76–81.

Shelley, J. J., Swearer, S. E., Adams, M., Dempster, T., Le Feuvre, M. C., Hammer, M. P., & Unmack, P. J. (2018). Cryptic biodiversity in the freshwater fishes of the Kimberley endemism hotspot, northwestern Australia. Molecular Phylogenetics and Evolution, 127, 843–858.

Stamatakis, A. 2014. RAxML version 8: a tool for phylogenetic analysis and post-analysis of large phylogenies. Bioinformatics, 30, 1312–1313.

Sunnucks, P., Morales, H. E., Lamb, A. M., Pavlova, A., & Greening, C. (2017). Integrative approaches for studying mitochondrial and nuclear genome co-evolution in oxidative phosphorylation. Frontiers in Genetics, 8, 25.

Townsend, C. R., & Crowl, T. A. (1991). Fragmented population structure in a native New Zealand fish: an effect of introduced brown trout? Oikos, 61, 347–354.

Unmack, P. J., Hammer, M. P., Adams, M., & Dowling, T. E. (2011). A phylogenetic analysis of pygmy perches (Teleostei: Percichthyidae) with an assessment of the major historical influences on aquatic biogeography in southern Australia. Systematic Biology, 60, 797–812.

Unmack, P. J., Sandoval-Castillo, J., Hammer, M. P., Adams, M., Raadik, T. A., & Beheregaray, L. B. (2017). Genome-wide SNPs resolve a key conflict between sequence and allozyme data to confirm another threatened candidate species of river blackfishes (Teleostei: Percichthyidae: Gadopsis). Molecular Phylogenetics and Evolution, 109, 415–420.

Veale, A. J., & Russello, M. A. (2017). Genomic changes associated with reproductive and migratory ecotypes in sockeye salmon (Oncorhynchus nerka). Genome Biology and Evolution, 9, 2921–2939.

Wallis, G. P., Cameron-Christie, S. R., Kennedy, H. L., Palmer, G., Sanders, T. R., & Winter, D. J. (2017). Interspecific hybridization causes long-term phylogenetic discordance between nuclear and mitochondrial genomes in freshwater fishes. Molecular Ecology, 26, 3116–3127.

Ward, R. D., Woodwark, M., & Skibinski, D. O. F. (1994). A comparison of genetic diversity levels in marine, freshwater, and anadromous fishes. Journal of Fish Biology, 44, 213–232.

Waters, J. M., & Burridge, C. P. (2016). Fine-scale habitat preferences influence within-river population connectivity: a case-study using two sympatric New Zealand Galaxias fish species. Freshwater Biology, 61, 51–56.

Waters, J. M., Burridge, C. P., & Craw, D. (2020b). River capture and freshwater biological evolution: A review of galaxiid fish vicariance. Diversity, 12, 216.

Waters, J. M., Craw, D., Burridge, C. P., Kennedy M., King, T. A., & Wallis, G. P. (2015b). Within-river genetic connectivity patterns reflect contrasting geomorphology. Journal of Biogeography, 42, 2452–2460.

Waters, J. M., Wallis, G.P., Burridge, C.P., Craw, D. (2015a). Geology shapes biogeography: Quaternary river-capture explains New Zealand’s biologically ‘composite’ Taieri River. Quaternary Science Reviews, 120, 47–56.

Waters, J. M., Craw, D., Youngson, J. H., & Wallis, G. P. (2001a). Genes meet geology: fish phylogeographic pattern reflects ancient, rather than modern, drainage connections. Evolution, 55, 1844–1851.

Waters J. M., Emerson, B. C., Arribas, P., & McCulloch, G. A. (2020a). Dispersal reduction: causes, genomic mechanisms, and evolutionary consequences. Trends in Ecology and Evolution, 35, 512–522.

Waters, J. M., Esa, Y. B., & Wallis, G. P. (2001b). Genetic and morphological evidence for reproductive isolation between sympatric populations of Galaxias (Teleostei: Galaxiidae) in South Island, New Zealand. Biological Journal of the Linnean Society, 73, 287–298.

Waters, J. M., Rowe, D. L., Burridge, C. P., & Wallis, G. P. (2010). Gene trees versus species trees: reassessing life-history evolution in a freshwater fish radiation. Systematic Biology, 59, 504–17.

Waters, J. M., Shirley, M., & Closs, G. P. (2002). Hydroelectric development and translocation of Galaxias brevipinnis: a cloud at the end of the tunnel? Canadian Journal of Fisheries and Aquatic Sciences, 59, 49–56.

Waters, J. M., & Wallis, G. P. (2001a). Cladogenesis and loss of the marine life-history phase in freshwater galaxiid fishes (Osmeriformes: Galaxiidae). Evolution, 55, 587–597.

Waters, J. M., & Wallis, G. P. (2001b). Mitochondrial DNA phylogenetics of the Galaxias vulgaris complex from South Island, New Zealand: rapid radiation of a species flock. Journal of Fish Biology, 58, 1166–1180.

Whiteley, A. R., Fitzpatrick, S. W., Funk, W. C., & Tallmon, D. A. (2015). Genetic rescue to the rescue. Trends in Ecology and Evolution 30:42–49.

Willis, S. C., Farias, I. P. & Ortí, G. (2014). Testing mitochondrial capture and deep coalescence in Amazonian cichlid fishes (Cichlidae: Cichla). Evolution, 68, 256–268.

